# Generating realistic null hypothesis of cancer mutational landscapes using SigProfilerSimulator

**DOI:** 10.1101/2020.02.13.948422

**Authors:** Erik N. Bergstrom, Mark Barnes, Iñigo Martincorena, Ludmil B. Alexandrov

## Abstract

Performing a statistical test requires a null hypothesis. In cancer genomics, a key challenge is the fast generation of accurate somatic mutational landscapes that can be used as a realistic null hypothesis for making biological discoveries. Here we present SigProfilerSimulator, a powerful tool that is capable of simulating the mutational landscapes of thousands of cancer genomes at different resolutions within seconds. Applying SigProfilerSimulator to 2,144 whole-genome sequenced cancers reveals: *(i)* that most doublet base substitutions are not due to two adjacent single base substitutions but likely occur as single genomic events; *(ii)* that an extended sequencing context of +/-2bp is required to more completely capture the patterns of substitution mutational signatures in human cancer; *(iii)* information on false-positive discovery rate of commonly used bioinformatics tools for detecting driver genes. SigProfilerSimulator’s breadth of features allows one to construct a tailored null hypothesis and use it for evaluating the accuracy of other bioinformatics tools or for downstream statistical analysis for biological discoveries.

## MAIN

Performing a statistical evaluation to determine whether an observation is seen by chance necessitates the construction of a null hypothesis corresponding with the expected default position. An observation is generally considered statistically significant if it reflects an unlikely outcome of the null hypothesis. In most practical applications, observations seen in less than 5% of outcomes from a null distribution are considered statistically significant.

Large-scale computational analyses of cancer genomes use background mutational models to evaluate driver mutations [1-6], mutational signatures [7], and topographical accumulation of somatic mutations [8]. In almost all cases, a null hypothesis model of the background mutation rate is implicitly incorporated into a bioinformatics tool [6, 9, 10] and used to report statistically significant results. Here we present SigProfilerSimulator, a computationally efficient bioinformatics tool for generating sample specific mutational landscapes that match the mutational signatures operative in each sample. SigProfilerSimulator provides a framework for generating a background mutational model for downstream statistical analyses and hypothesis testing. The tool supports generation of simulated single base substitutions (SBSs), small insertions and deletions (IDs), and doublet base substitutions (DBSs) while maintaining their patterns at different resolutions. SigProfilerSimulator is available as both a Python and an R package, provides support for commonly used data formats, and is extensively documented. To demonstrate the wide applicability of SigProfilerSimulator, we illustrate its basic functionality using a single cancer genome and then apply the tool to 2,144 whole-genome sequenced cancers and to 1,024 whole-exome sequenced breast cancers to address three different questions in cancer genomics.

The mutational pattern of a cancer genome can be described using distinct classification schemes reflecting the activity of mutational processes at different resolutions [11]. For example, single base substitutions can be described using only the mutated base-pair (6 possible mutational channels; known as SBS-6 classification), or the mutated base-pair with +/-1bp context (SBS-96), or the mutated base-pair with +/-2bp context (SBS-1536), etc. [12]. Each of these classifications can be subsequently elaborated by considering additional features, e.g., transcriptional strand bias [11, 12]. By preserving the pattern of mutations at a preselected resolution, SigProfilerSimulator converts a set of real somatic mutations from a cancer genome into another set of randomly generated somatic mutations (**Fig. 1*A***). Maintaining the mutational pattern provides an assurance that the same mutational processes are observed in both the real and the simulated cancer genome. By default, the tool projects these mutations as statistically independent events onto each chromosome by proportionately assigning mutations based on both the chromosome’s length and the observed rate of each mutational channel in each chromosome of a preselected reference genome. The tool also provides a variety of custom options for simulating mutations, including: *(i)* gender of the sample allowing appropriate incorporation of sex chromosomes; *(ii)* transcriptional strand bias allowing accurate distribution of mutations to account for the activity of transcription-coupled nucleotide excision repair; *(iii)* considering mutations as dependent sequential events, i.e., each mutation updates the observed rate of a mutational channel for a preselected reference genome; *(iv)* preserving mutational burden and mutational patterns for each chromosome; *(v)* exome simulations that generate mutations only in the coding regions of the genome; and several other options. With this collection of features, one can easily tailor an appropriate background mutational model for testing different biological hypotheses or for evaluating existing bioinformatics tools. Importantly, SigProfilerSimulator is computationally efficient. For example, the tool can simulate ∼37 million somatic mutations found in the 2,144 whole-genome sequenced cancers generated by Pan-cancer Analysis of Whole Genomes (PCAWG) initiative [13] within 90 seconds.

**Figure 1.**
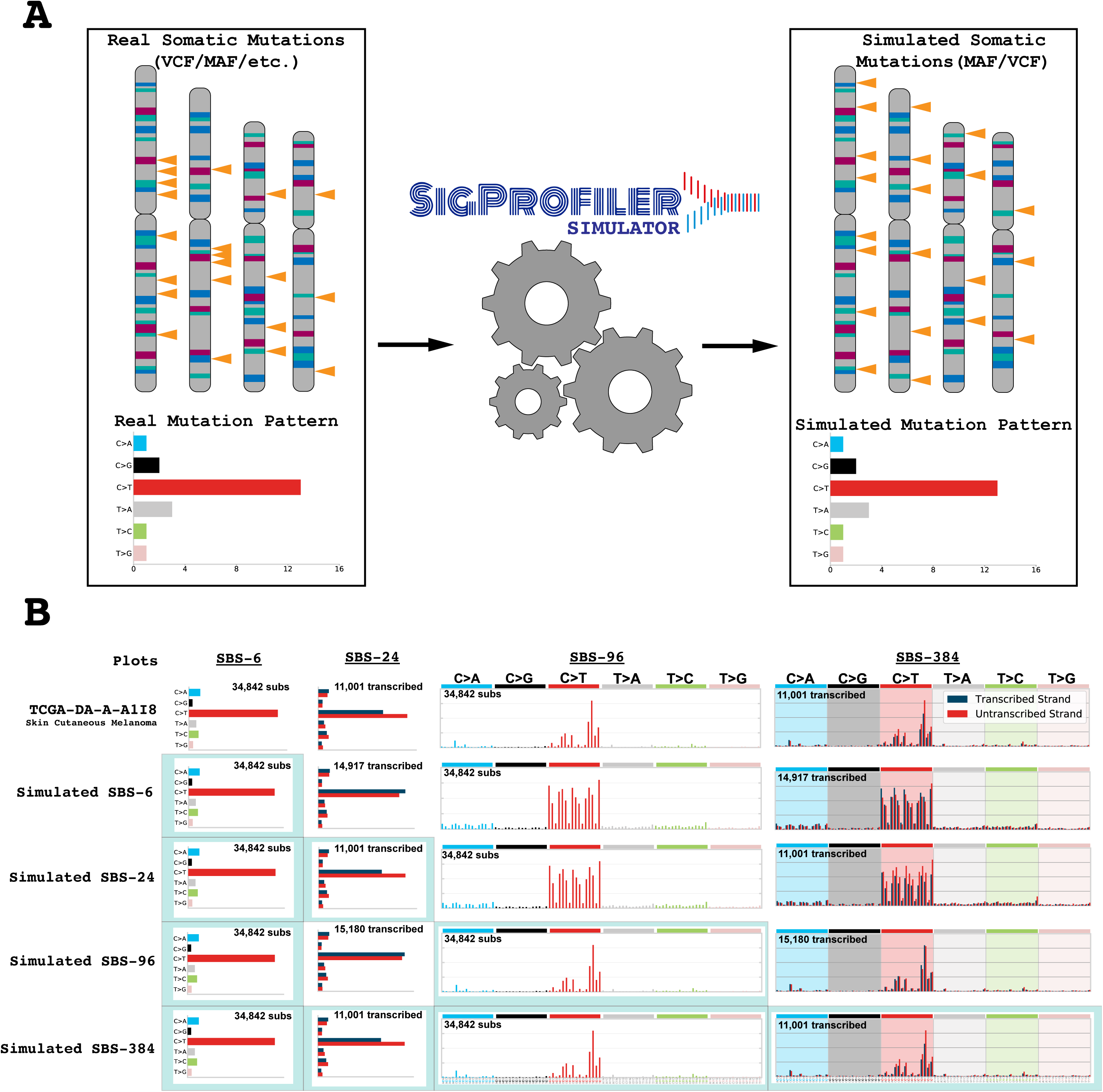
High-level overview illustrating the functionality of SigProfilerSimulator. ***A)*** Schematic depiction of SigProfilerSimulator’s general functionality. The tool transforms the real somatic mutational catalog of a cancer genome into a simulated mutational catalog, while maintaining the mutational burden and the mutational pattern at a preselected resolution. ***B)*** Comparing the simulated catalogues of a single cancer genome at different resolutions. Adding additional sequence context to a simulation creates a more specific and complex mutational model. Similarly, one can preserve the transcriptional strand bias (SBS-24, SBS-384, and SBS-6144) to incorporate additional specificity. Simulating a more complex classification of the data results in matching catalogs for all collapsed versions of the higher matrix (i.e., simulating SBS-384 ensures that the SBS-6, SBS-24, and SBS-96 simulated catalogs match the original data).

To illustrate several of SigProfilerSimulator’s features, we provide a detailed visualization for a single TCGA melanoma sample: TCGA-DA-A-A1I8. Simulating TCGA-DA-A-A1I8 using the SBS-6 classification maintains the original sample’s pattern for the six possible types of single base mutations, however, it also results in completely different patterns for classifications at higher resolutions (**Fig. 1*B***). Simulating an extended sequence context (SBS-96) results in a perfect match with the original landscape when including +/-1 adjacent bases; however, it does not reflect the transcriptional strand bias observed in the sample (**Fig. 1*B***). As such, one can further elaborate these simulations by incorporating transcriptional strand bias (**Fig. 1*B***), by considering +/-2 adjacent bases (**Supplemental Fig. 1**), or by preserving the mutational burden and mutational patterns on each chromosome (**Supplemental Fig. 1**). Each of these simulations can be subsequently used to test different hypotheses. To demonstrate this functionality, we applied SigProfilerSimulator to three questions in cancer genomics.

First, we used simulations to evaluate whether doublet base substitutions (e.g., CC:GG>TT:AA mutations) are two subsequent single base substitutions occurring simply by chance in adjacent genomic positions. We constructed a null hypothesis by applying the tool to the 2,144 PCAWG cancer genomes. Simulations were performed considering SBSs as both statistically independent events (non-updating – simulating with replacement; each mutation has no effect on the observed rate of mutational channels) and dependent events (updating – simulating without replacement; each mutation updates the observed rate of mutational channels). Each sample was simulated 1,000 times providing a distribution of DINUCs. After simulating the SBS-96 context for each PCAWG sample, we examined the number of single base substitutions occurring next to one another simply by chance. For example, in the sample SP99325 (LIRI), we observed on average approximately 23 pairs of adjacent SBSs when considering mutations as statistically independent events and 14 pairs of adjacent SBSs when considering mutations as dependent events (**Fig. 2*A***). In contrast, the actual sample contains 303 doublet base substitutions indicating a 22-fold and a 13-fold enrichment compared to the null hypothesis, respectively. The results indicate that it is highly unlikely that the majority of observed doublet base substitutions in SP99325 are the result of two adjacent SBS events. Applying the same approach to all PCAWG samples reveals between 10- and 1000-fold increase of the real number of DBSs compared to simulated data (**Fig. 2*B*; Supplemental Fig. 2**). These results confirm the believe that the vast majority of doublet base substitutions in human cancer are not due to adjacent single base substitutions. Rather, doublet base substitutions are likely due either to single genomic events or to higher mutagenic propensities of certain regions of the human genome.

**Figure 2.**
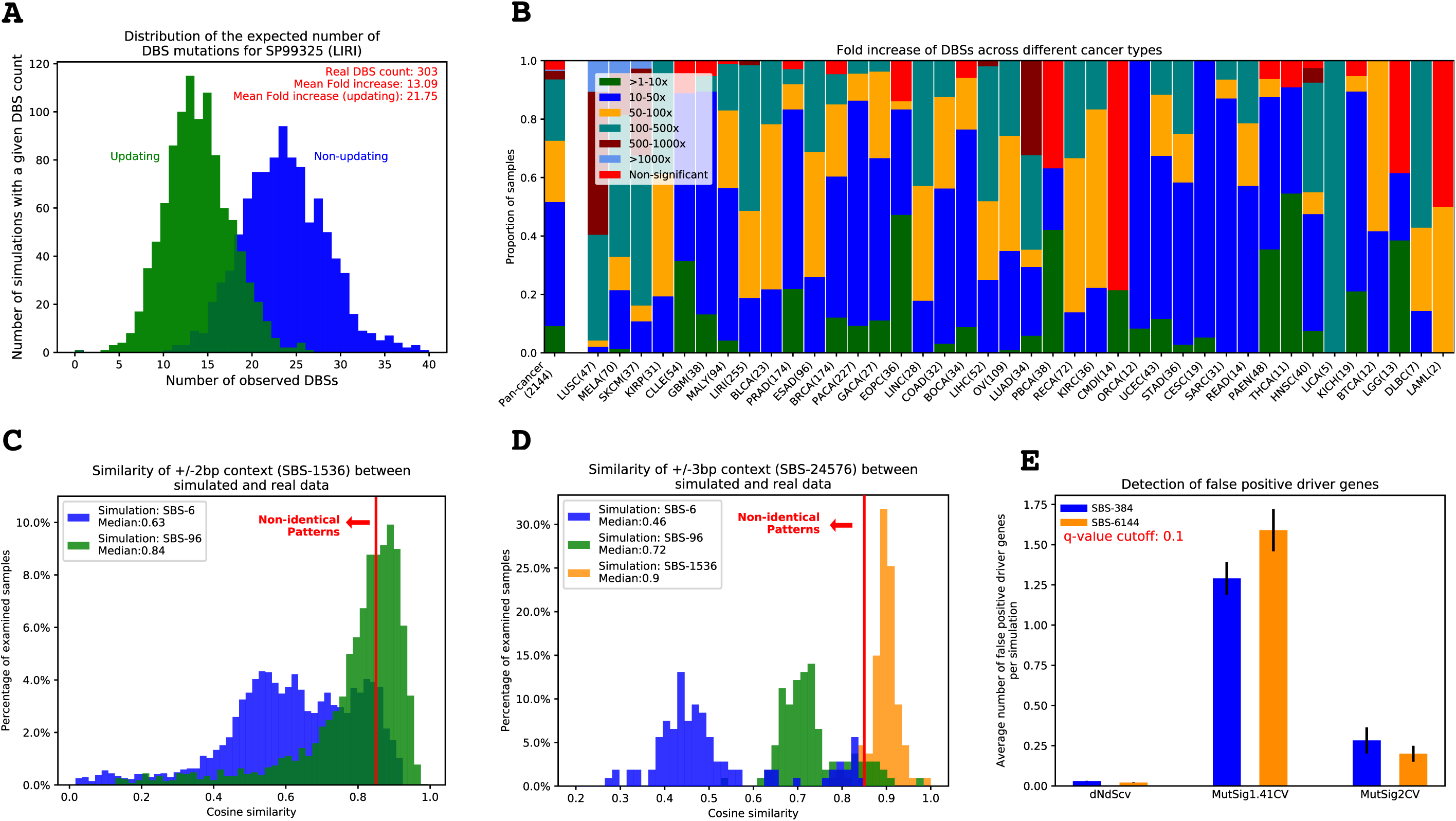
Applying SigProfilerSimulator to three distinct cancer genomics problems. ***A)*** Distribution of the expected number of doublet base substitutions (DBSs) due to the adjacent single base substitutions (SBSs) observed by chance for the PCAWG sample SP99325. The distributions represent the results from 1,000 simulations of the mutational pattern of SP99325 treating mutations as statistically independent events (blue) and 1,000 simulations of the mutational pattern of SP99325 treating mutations as dependent events. ***B)*** The fold increase of DBSs observed in the original PCAWG samples and the average number of DBSs observed in our simulations. The mutational pattern of each sample was generated 1,000 times considering somatic mutations as statistically independent events. ***C)*** Comparing the similarities of mutational patterns at +/-2bp context (SBS-1536) between real and simulated PCAWG samples. Simulations were performed at SBS-6 and SBS-96 resolutions. ***D)*** Comparing the similarities of mutational patterns at +/-3bp context (SBS-24576) between real and simulated PCAWG samples. Simulations were performed at SBS-6, SBS-96, and SBS-1536 resolutions. ***E)*** Evaluating the false-positive rates of MutSigCV1.41, MutSigCV2, and dNdScv driver detection tools using SigProfilerSimulator. All TCGA breast cancer WES samples were simulated 100 times and examined for driver mutations using both MutSigCV and dNdScv. The average number of significant driver genes are plotted using a recommended q-value cutoff of 0.10.

Second, we evaluated whether incorporating additional sequence context 5’ and 3’ of single base substitutions increases the specificity of the mutational patterns observed in cancer genomes [12]. Here, we considered two mutational patterns to be the same if their cosine similarity is more than 0.85 (Methods). Specifically, we simulated the PCAWG dataset at different resolutions (*viz.*, SBS-6, SBS-96, and SBS-1536) and compared them to the patterns of mutations observed in the real samples (**Fig. 2*C*** & **2*D***). Comparing the +/-2bp context of data simulated using SBS-6 to the +/-2bp context of the real data demonstrated that for almost all samples the SBS-6 simulations do not capture the +/-2bp context as 91% of samples exhibited a cosine similarity below 0.85. Similarly, only half of the samples simulated using SBS-96 had consistent +/-2bp context when compared to the real data (44% below 0.85; **Fig. 2*C***). This demonstrates that the mutational patterns of the examined cancer genomes exhibit additional specificity for +/-2bp adjacent to single base substitutions. In contrast, comparing the +/-3bp context of data simulated using SBS-1536 demonstrated that the +/-2bp context captures the patterns observed at +/-3bp for almost all samples (only 6.5% of samples below 0.85; **Fig. 2*D***). Overall, these results suggest that the SBS-1536 classification is necessary to capture additional information for a set of signatures beyond SBS-6 and SBS-96. Moreover, extending this classification to +/-3bp (SBS-24576) is largely not necessary as the SBS-1536 classification already captures the patterns of +/-3bp for majority of the examined cancer samples.

Third, we evaluated the false-positive rates of tools commonly used for discovery of cancer driver genes. More specifically, we simulated the somatic mutations observed in the 1,024 whole-exome sequenced breast cancers reported in the TCGA MC3 release [14]. The simulations were repeated 100 times and each of these 100 repetitions was analyzed for driver genes using MutSigCV1.41 and MutSigCV2 [6] as well as dNdScv [10]. In principle, since SigProfilerSimulator randomly shuffles somatic mutations, one would not expect to find any genes under selection. However, each of the tools found significantly mutated genes within the simulations using the recommended cutoff threshold of q-value < 0.10 (**Fig. 2*E***). On average MutSig1.41CV found between 1.3 and 1.6 false-positive driver genes per simulation when examining data generated using the SBS-384 and SBS-6144 mutational classifications, respectively. In contrast, MutSig2CV found between 0.3 and 0.2 false-positive driver genes per simulation using SBS-384 and SBS-6144, respectively. Lastly, dNdScv found between 0.03 and 0.02 false-positive driver genes per simulation using SBS-384 and SBS-6144, respectively. Note that by chance, when using a q-value cutoff of 0.1, one would expect to observe less than 0.1 false-positive driver genes per simulation. Lowering the threshold for statistical significance to 0.01 eliminates all false-positive results from dNdScv and MutSig2CV but not for MutSig1.41CV.

Increasingly, there is a need to develop reliable background models of cancer mutational landscapes to allow downstream statistical analysis for biological discoveries. Currently, to the best of our knowledge, there is no tool that allows explicitly simulating accurate background mutational landscapes. This report presents SigProfilerSimulator, a method that allows fast generation of mutational landscapes at different resolutions. As demonstrated by our analyses, SigProfilerSimulator can be used to evaluate the accuracy of other bioinformatics tools or it can be leveraged for making novel discoveries. SigProfilerSimulator’s breadth of features allows one to construct a tailored null hypothesis of mutational landscapes and to identify significance levels of the subsequent results. Overall, SigProfilerSimulator will be a useful tool for any researcher that performs statistical analysis based on mutational data derived from the sequencing of cancer or normal somatic tissues.

## METHODS

### Tool Implementation

SigProfilerSimulator is developed as a computationally efficient Python package and it is available for installation through PyPI. Further, an R-wrapper is available through GitHub. The tool leverages a PCG random number generator that provides a simple, fast, and space-efficient algorithm for generating random numbers with high statistical quality [15]. The tool uses a Monte Carlo approach for randomly generating somatic mutations while considering the observed frequency of a preselected reference genome. More specifically, SigProfilerSimulator randomly shuffles mutations by using the precomputed observed rates of mutational channels in a reference genome. The tool works in unison with SigProfilerMatrixGenerator [11] to first classify a catalog of somatic mutations prior to simulating it. The final mutational catalog is outputted into commonly used mutation data formats including mutation annotation format (MAF) files and variant annotation format (VCF) files. SigProfilerSimulator is freely available and has been extensively documented.

*Python code:* https://github.com/AlexandrovLab/SigProfilerSimulator

*R wrapper:* https://github.com/AlexandrovLab/SigProfilerSimulatorR

*Documentation:* https://osf.io/usxjz/wiki/home/

### Computational Benchmarking

The computational efficiency of SigProfilerSimulator was benchmarked by simulating the freely available PCAWG dataset, consisting of 2,144 samples with 36,876,213 single base substitutions, for a single iteration using the default parameters. Simulating the complete dataset took approximately 90 seconds. Simulations were performed on a dedicated computational node with a dual Intel® Xeon® Gold 6132 Processors (19.25M Cache, 2.60 GHz) and 192GB of shared DDR4-2666 RAM.

### Analysis of Doublet Base Substitutions

We simulated the PCAWG dataset using the SBS-96 classification. Each simulation was performed 1,000 times considering mutations as both statistically independent events (non-updating; each mutation has no effect on the observed rate of mutational channels) and dependent events (updating; each mutation updates the observed rate of mutational channels). To calculate the number of DBS mutations occurring by chance in each sample, we generated the mutational catalogs for DBS-78 using SigProfilerMatrixGenerator [11]. The resulting counts for DBSs were used to plot the distributions of the expected number of DBSs due to two adjacent SBSs happening purely by chance. The fold change was calculated by dividing the mean DBS count observed across the simulations by the total number of DBSs found in the original sample. Derivation of q-values was performed by applying the Benjamini and Hochberg false discovery rate correction to p-values calculated using z-tests based on the DBS distributions found in the simulations and the numbers of DBSs observed in the real data.

### Sequence Context Analysis for Mutational Signatures

The PCAWG dataset was simulated using the SBS-6, SBS-96, and SBS-1536 classifications while ensuring the respective mutational patterns and mutational burdens on each chromosome match the ones observed in the real data. SigProfilerMatrixGenerator was used to derive the mutational vectors for each sample including vectors incorporating three bases 5’ and three bases 3’ of each mutation, resulting in a classification with 24,576 mutational channels. To avoid comparisons of sparse binary vectors, only samples that had at least 2 mutations per mutational channel were included in the comparative analyses. The simulated and real mutational patterns of a cancer genome were considered the same if their cosine similarity was at least 0.85. Note that the average cosine similarity between two random nonnegative vectors is 0.75 (**Supplemental Fig. 3**). The chance of two nonnegative vectors with 1,536 mutational channels or 24,576 mutational channels to have a similarity of 0.85 simply by chance less than 10^−6^ (**Supplemental Fig. 3**).

### Benchmarking False-Positive Driver Genes Detected by MutSigCV and dNdScv

All whole-exome sequenced breast cancer samples part of the TCGA MC3 release were simulated using SBS-384 and SBS-6144 contexts while maintaining the mutational burden on each chromosome. As recommended [10], 23 samples with more than 500 exonic mutations were excluded from the analysis. Each simulation was repeated 100 times with different random seeds. The variant annotation predictor [16] was used to annotate simulated mutations with the appropriate gene name for compatibility with MutSigCV1.41 and MutSigCV2 [6]. We ran MutSigCV1.41 and MutSigCV2 using the recommended default parameters in conjunction with the genome reference sequence for hg19, mutation dictionary file, exome coverage file, and gene covariates file as found at https://software.broadinstitute.org/cancer/cga/mutsig_run. We ran dNdScv [10] using the default library parameters and filtered out the significant genes using the recommended q-value cutoff of less than 0.10.

## DECLARATIONS

### Ethics approval and consent to participate

Not applicable.

### Consent for publication

Not applicable.

### Availability of data and materials

No novel data were generated as part of this study. All source code is freely available and can be downloaded from the links below.

*Python code:* https://github.com/AlexandrovLab/SigProfilerSimulator

*R wrapper:* https://github.com/AlexandrovLab/SigProfilerSimulatorR

*Documentation:* https://osf.io/usxjz/wiki/home/

### Author Contributions

E.N.B. developed the Python and R code and wrote the manuscript. M.B. tested and documented the code. I. M. provided advice for both driver gene analysis and the interpretation of the results. L.B.A. supervised the overall development of the code and writing of the manuscript. All authors read the manuscript.

## Acknowledgements

The authors thank M. Lawrence for sharing the newest code for MutSig2CV. The authors also thank M. Zhivagui for designing the SigProfiler logo.

## SUPPLEMENTARY INFORMATION

**Supplementary Figure 1. Example of an additional resolution for simulating mutational patterns supported by SigProfilerSimulator**. The example illustrates the resulting patterns when maintaining the mutational burden on each chromosome and when only relying on proportionate allocation based upon the nucleotide context distribution of the reference genome. Comparison is provided for a single breast cancer sample simulated at an SBS-1536 resolution.

**Supplementary Figure 2. Evaluating the expected rates of DBSs for mutations simulated as dependent events.** The fold increase of DBSs observed in the original PCAWG samples and the average number of DBSs observed in our simulations. The mutational pattern of each sample was generated 1,000 times considering somatic mutations as dependent events.

**Supplementary Figure 3. Evaluating the average similarity of random nonnegative vectors. *A)*** Comparing the cosine similarities amongst 10,000 randomly generated nonnegative vectors, where each vector has 1,536 mutational channels. ***B)*** Comparing the cosine similarities amongst 10,000 randomly generated nonnegative vectors, where each vector has 24,576 mutational channels.

