## Supplemental Figures for "Generating realistic null hypothesis of cancer mutational landscapes using SigProfilerSimulator"

### Chromosome 10

Supplementary Figure 1

**PD6413a**
**Breast cancer**

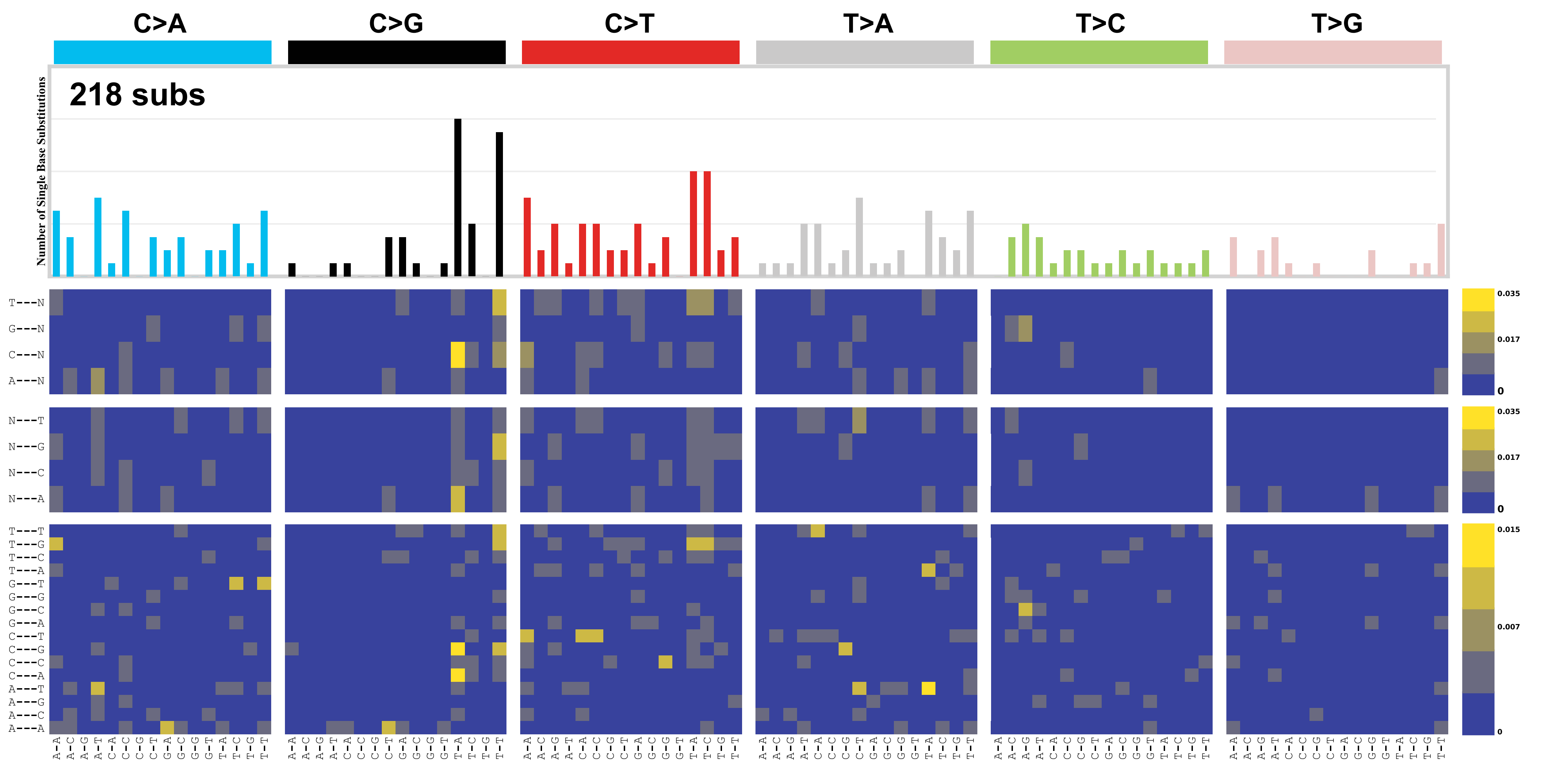


**Simulated SBS-1536**

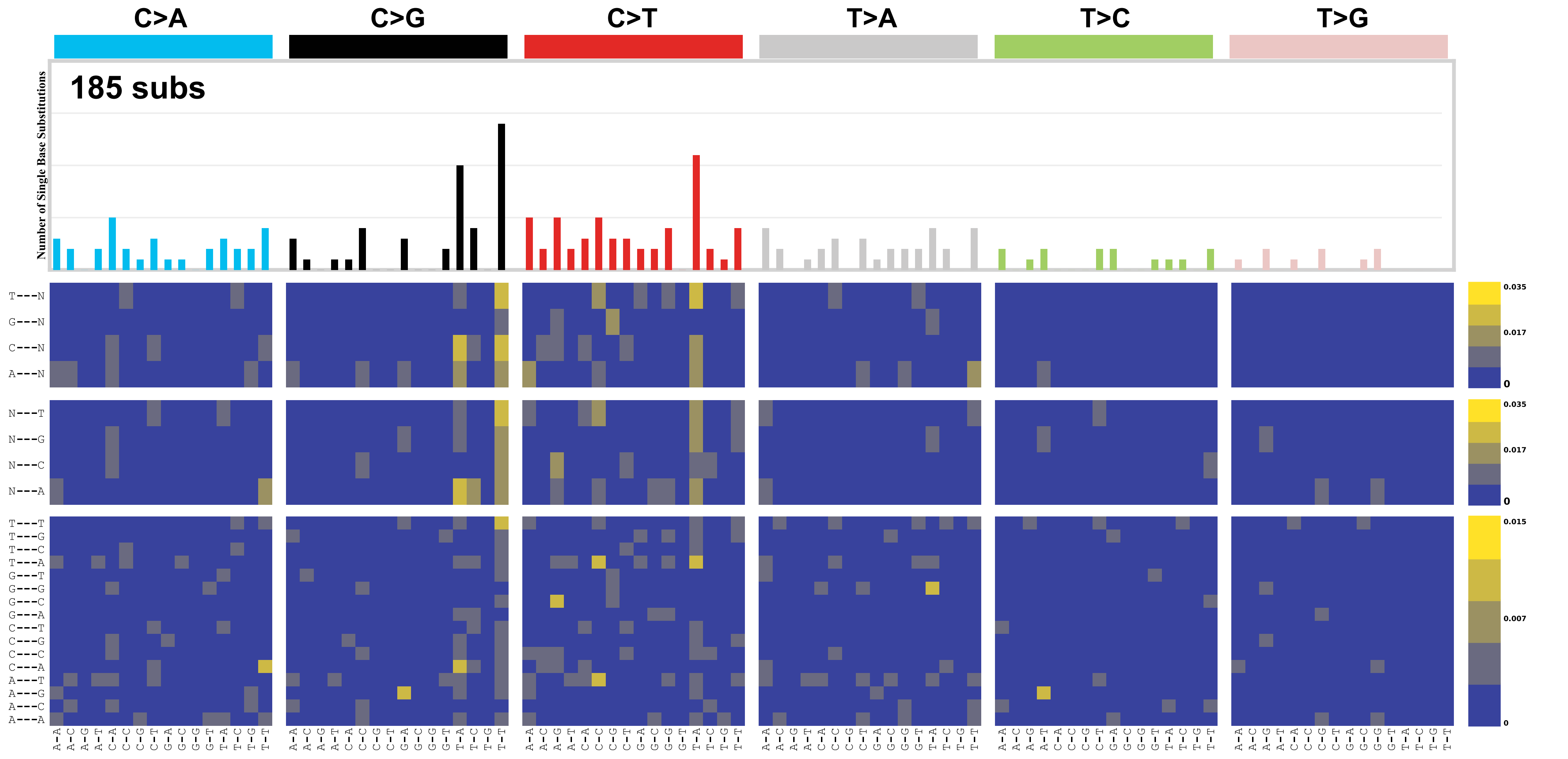


**Simulated SBS-1536 per chromosome**

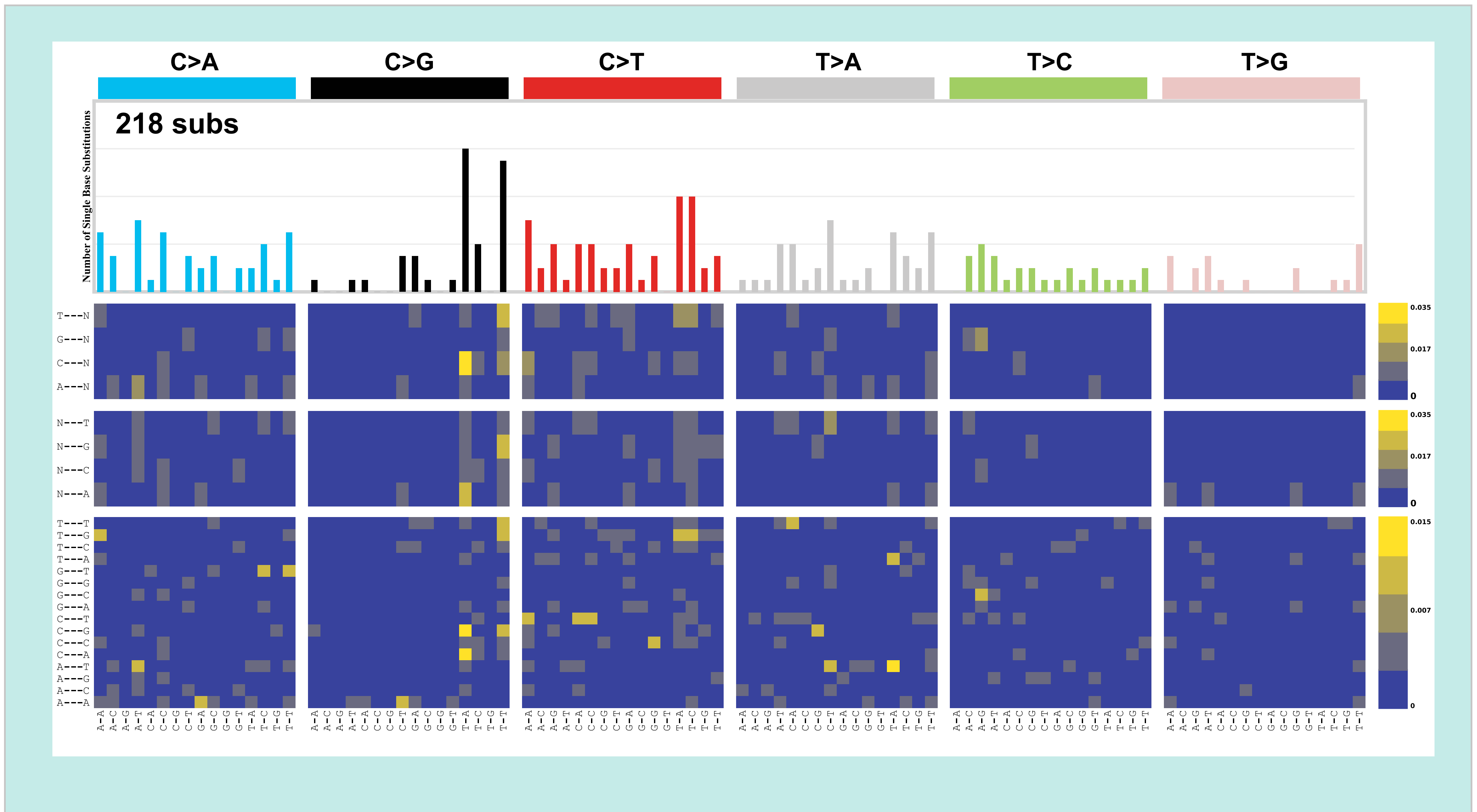

**Supplementary Figure 2**

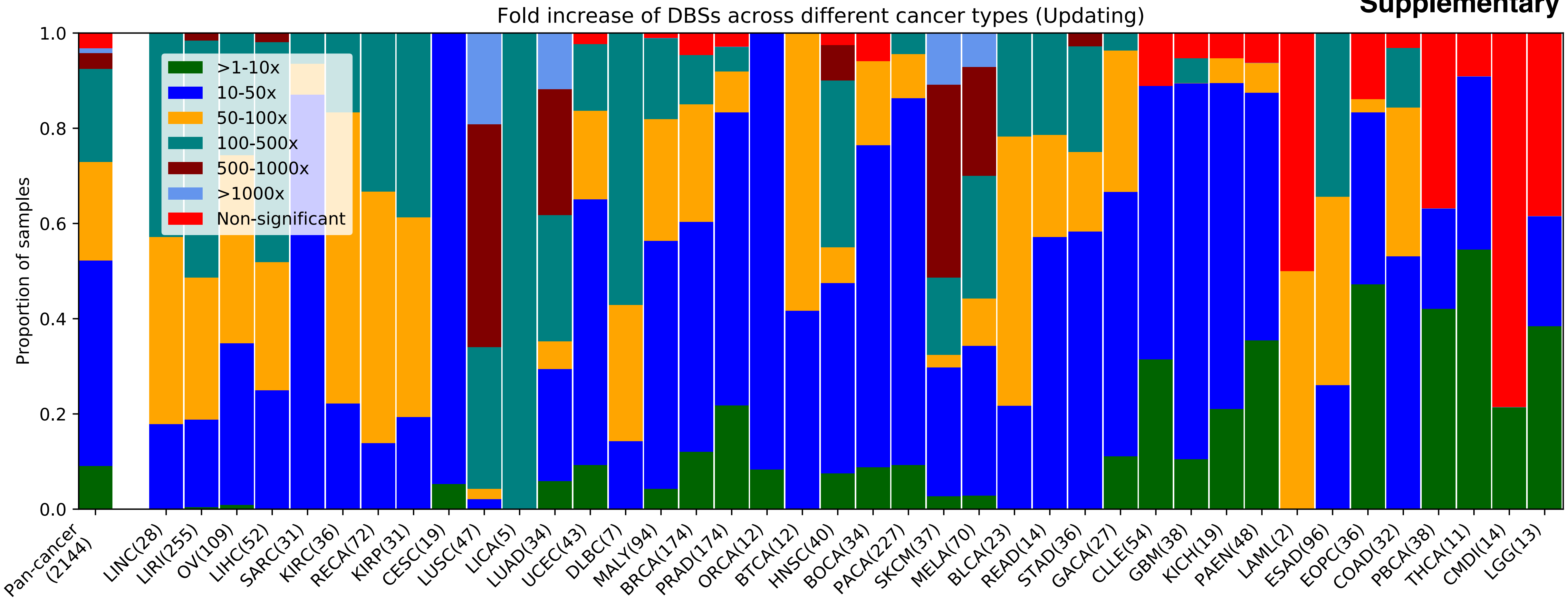

### Supplementary Figure 3

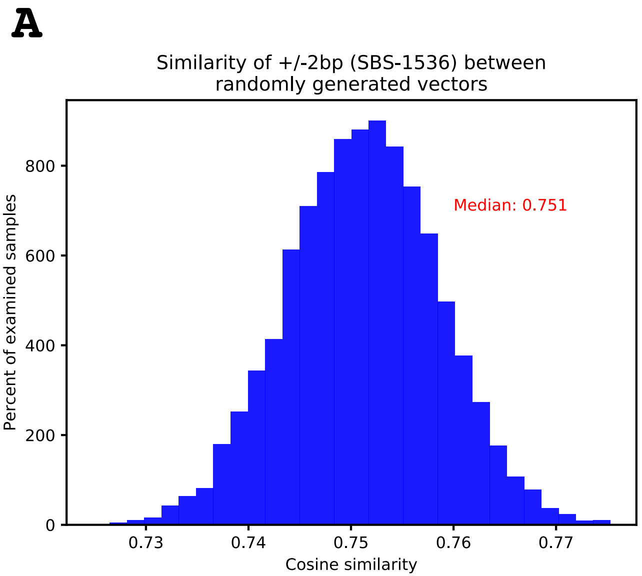


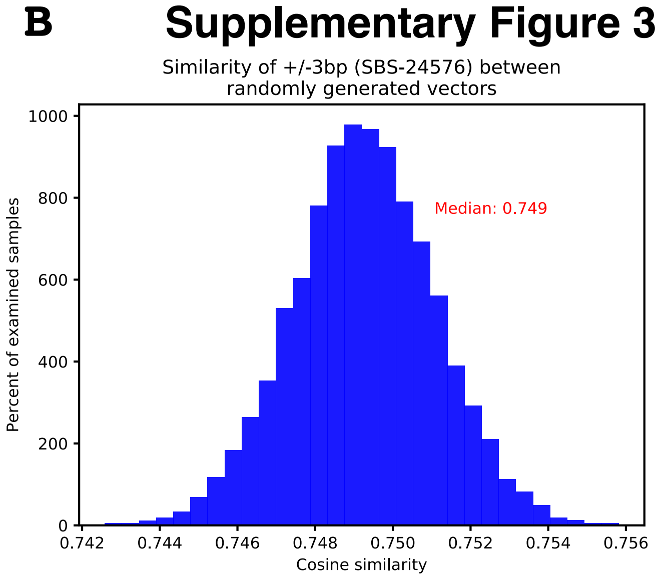
